# Cholinergic modulation of motor sequence learning

**DOI:** 10.1101/2023.10.11.561645

**Authors:** Angela Voegtle, Catharina Mohrbutter, Jonathan Hils, Steve Schulz, Alexander Weuthen, Uwe Brämer, Markus Ullsperger, Catherine M. Sweeney-Reed

## Abstract

The cholinergic system plays a key role in motor function, but whether pharmacological modulation of cholinergic activity affects motor sequence learning is unknown. The acetylcholine receptor antagonist biperiden, an established treatment in movement disorders, reduces attentional modulation, but whether it influences motor sequence learning is not clear. Using a randomized, double-blind placebo-controlled crossover design, we tested thirty healthy young participants and show that biperiden impairs production of sequential finger movements following a fixed but not a random sequence. A similar interaction was observed in widespread oscillatory broadband power changes (4-25 Hz) in the motor sequence learning network after receiving biperiden, with greater power in the theta, alpha, and beta bands over ipsilateral motor and bilateral parietal–occipital areas. The reduced theta power during a fixed compared to random sequence, likely reflecting disengagement of top-down attention to sensory processes, was disrupted by biperiden. The alpha synchronization during learned sequences, reflecting sensory gating and lower visuospatial attention requirements for the learned, compared with visuomotor responses to a random sequence, was greater after biperiden, potentially reflecting excessive visuospatial attention reduction following biperiden, also affecting visuomotor responding required to enable sequence learning. Beta oscillations facilitate sequence learning by integrating visual and somatosensory inputs, stabilizing learned sequences, and promoting prediction of the next stimulus. The beta synchronization after biperiden fits with a disruption of the selective visuospatial attention enhancement associated with initial sequence learning. These findings highlight the role of cholinergic processes in motor sequence learning.

## Introduction

Motor sequence learning, a form of procedural memory, is critical to performance of regular daily activities, such as riding a bicycle. The neurotransmitter acetylcholine (ACh), which plays a key role in movement disorders such as Parkinsons’s disease and dystonia (Brocks, 1999; Eskow Jaunarajs et al., 2015), is also involved in motor learning (Conner et al., 2003; Conner et al., 2010). The ACh receptor antagonist biperiden, an established treatment for patients with Parkinson’s disease and dystonia (Bezchlibnyk-Butler & Remington, 1994; Brocks, 1999; Kawabata & Katsuno, 2021), has been shown to impair episodic, verbal, and working memory (Borghans et al., 2020; Guthrie et al., 2000; Klinkenberg & Blokland, 2011; Sambeth et al., 2015; Wezenberg et al., 2005). However, the influence of ACh inhibition on motor functioning is less clear. While some studies have shown no impact on general motor execution (Borghans et al., 2020; Guthrie et al., 2000), others have identified impairments on sensorimotor (Klinkenberg & Blokland, 2011; Wezenberg et al., 2005), visuo-spatial (Linstow Roloff et al., 2007; Sambeth et al., 2015; Wezenberg et al., 2005), and procedural memory tasks (Rasch et al., 2009). Biperiden has a high affinity for the muscarinic M1 receptor (Bolden et al., 1992), which is primarily located in the hippocampus and the cerebral cortex, but also in the cerebellum, striatum, and thalamus (Levey, 1993). Neuroimaging, anatomical, and lesion studies have identified a cortical–subcortical network underpinning procedural learning, engaging a frontal–parietal network (Eliassen et al., 2001; Hardwick et al., 2013; Müller et al., 2002; Rivera-Urbina et al., 2022; Tzvi et al., 2016) and the motor cortex, cerebellum, basal ganglia, and ventral intermediate nucleus of the thalamus (Hardwick et al., 2013; Roy et al., 2019; Terzic et al., 2022; Voegtle et al., 2022; Voegtle et al., 2023). The presence of M1 receptors in the network underpinning procedural learning suggests a potential impact of biperiden on motor learning. It is unknown, however, whether motor sequence learning is impaired through ACh inhibition. Patients with movement disorders including Parkinson’s disease and essential tremor can have impaired motor sequence learning (Meissner et al., 2018, 2019; Voegtle et al., 2023). Pharmacological modulation of brain ACh functioning therefore has important implications for treatment of patients suffering from movement disorders.

Motor sequence learning is accompanied by oscillatory spectral power modulation in the brain structures comprising the motor learning network. In addition to movement generation, beta (13-30 Hz) oscillations are considered to promote cortical plasticity (Pollok et al., 2014) and are altered in movement disorders (Meissner et al., 2018, 2019; Voegtle et al., 2023). Changes in alpha (8-12 Hz) spectral power have been associated with reduced cognitive control as a sequence becomes automatic (Pollok et al., 2014; Tzvi et al., 2016; Voegtle et al., 2023; Zhuang et al., 1997), and theta (4-8 Hz) oscillations may reflect attentional demands (Clayton et al., 2015; Lum et al., 2022; Meissner et al., 2018; Sauseng et al., 2007; Tzvi et al., 2016). Cholinergic modulations of alpha and beta oscillations are linked to spatial attention and sensory processing (Bauer et al., 2012; Knott et al., 1997; Neufeld et al., 1994; Osipova et al., 2003; Sannita et al., 1987), suggesting that oscillatory modulation could provide a window onto the effects of ACh on motor learning.

Here we assessed the impact of biperiden on motor sequence learning and its neural correlates in healthy participants using a randomized, double-blind, placebo-controlled study design, employing the serial reaction time test (SRTT), a well-established approach to evaluation of motor sequence learning (Nissen & Bullemer, 1987; Terzic et al., 2022; Voegtle et al., 2022). We hypothesized that motor sequence learning would be specifically impaired by biperiden, with modulation of oscillatory spectral power in the motor sequence learning network.

## Material & Methods

### Participants

Thirty healthy male adults aged 18–30 years were recruited through the Psychology Institute of the Otto von Guericke University, Magdeburg. The medical supervisor (UB, CMSR) screened volunteers for current or history of illnesses (cardiac, neurological, psychiatric, pulmonary, renal, and gastrointestinal), which were exclusion criteria, as were nicotine consumption, antihistamines taken on the same day, and intolerance to any of the components of the medication used (especially lactose or galactose). The Local Ethics Committee of the Otto von Guericke University, Magdeburg granted ethical approval. All participants provided informed, written consent before study inclusion, in accordance with the Declaration of Helsinki, and were informed of their right to cease participation at any time without providing reasons.

### Study design and treatment

A double-blind placebo-controlled crossover study design was applied, in which biperiden 4 mg and placebo were balanced over two sessions, at least 6 days apart. Participant body mass index (BMI) was recorded. Half of the participants received biperiden in the first testing session and half the placebo, in a counterbalanced order. The medication was in tablet form and provided by a local pharmacy identically for both preparations. On average the SRTT was performed 169 minutes after pharmacological intervention following different paradigms, which formed part of other studies. Before the SRTT, participants completed a German translation of a fatigue questionnaire containing 18 items and a scale with values between 1 and 10 (Lee et al., 1991). The scores for reversed-scale items were inverted and summed scores compared between conditions.

### Serial reaction time task

Four squares were shown on a computer screen, with stimulus–response mapping to the second to fifth fingers of the right hand. The squares turned red in an alternating fixed, repeated 12-item sequence (1-3-2-1-4-1-2-3-1-3-2-4) or in a random order, and the participants were asked to press the corresponding buttons as quickly and accurately as possible (Figure 1a). The stimuli were presented for 500 ms and the inter-stimulus interval was fixed at 1200 ms, independent of response times. Both sessions consisted of 3 blocks, each containing 288 trials. The 288 trials alternated between four cycles in which the stimuli followed a fixed sequence and four cycles in which the stimuli were selected at random, always starting with the sequential cycles (Figure 1b). At the end of each block, participants were asked whether they had noticed a repeated sequence, and if so to produce it.

**Figure 1.**
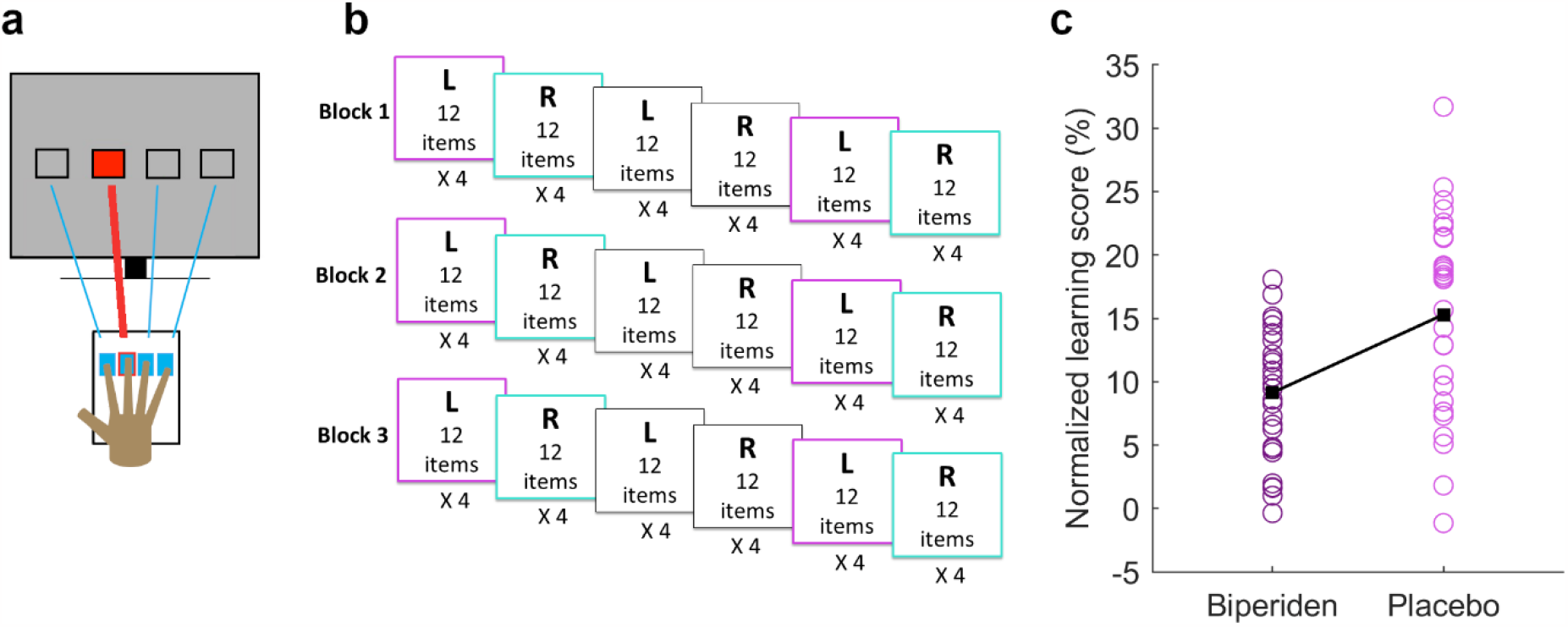
Serial reaction time task. **a**. The positions on the computer screen corresponding with the button positions. **b**. Three task blocks each comprised three runs of four fixed 12-item sequences, alternating with three runs of random sequences of 48 items. **c**. Individual participant normalized learning scores during the Biperiden and Placebo conditions.

### EEG recording and preprocessing

EEG was recorded with Brain Vision Recorder (Brain Products, Gilching, Germany) software using a 64-electrode equidistance EEG cap (Brain Products, Gilching, Germany) in a shielded chamber, with an electrode between FCz and aF3 set as the ground and electrode CPz as the reference. The sampling rate was 1000 Hz. Offline data processing was conducted with MATLAB R2018a (MathWorks, Natick, MA, USA) and Fieldtrip (version 20180826) (Oostenveld et al., 2011).

Raw data files were resampled to 500 Hz, band-pass filtered from 1 Hz to 100 Hz, and notch-filtered for 50 Hz line noise. The data were epoched from -1400-2600 ms around stimulus onset, with a 200 ms pre-stimulus baseline. Red highlighting of a square was considered a stimulus. Bad channels were removed from the data based on visual inspection. Independent component analysis was performed to eliminate eye artifacts. Afterwards, all epochs with values exceeding ±100 μV were excluded. Previously removed bad channels were replaced using spherical spline interpolation, and surface Laplacians were calculated. Time–frequency decomposition was performed through convolution with five-cycle complex Morlet wavelets for the frequencies 2-40 Hz from -200 ms to 1200 ms, normalizing the data with the 200 ms pre-stimulus. Event-related spectral perturbations (ERSPs) were determined for each participant by averaging over the trials with correct responses, separately for both sequence types (Learned, Random) and conditions (Biperiden, Placebo).

### Statistical analysis

Statistical analysis was performed using IBM SPSS Statistics 23 (IBM, Armonk, NY, USA), with MATLAB R2023b for visualization (MathWorks, Natick, MA, USA). ANOVAs and two-sided paired T-tests were applied if variables were normally distributed according to the Shapiro-Wilks test. If not, paired Wilcoxon rank sum tests were used.

Normalized learning scores ((mean RT to random - mean RT to learned)/mean RT to random)) were calculated for each individual based on the mean RTs for each *Condition* (Biperiden, Placebo).

A three-way repeated measures ANOVA was applied to the changes in RTs over each block with the within-subject factors *Condition* (Biperiden, Placebo), *Sequence type* (Learned, Random), and *Time* (Block 1, Block 2, Block 3), and the covariate *BMI*. Post hoc pairwise comparisons were Bonferroni-corrected. Means in ms and 95% confidence intervals are provided. Examining effects in the biperiden condition, a two-way repeated measures ANOVA was applied to the RT changes (collapsed over *Time*) with the within-subject factor *Sequence type*, the between-subject factor *Session order* (Biperiden first, Placebo first), and the covariates *BMI* and *Time after biperiden given*.

### Cluster-based permutation tests

To identify effects of biperiden on oscillatory spectral power, we performed three non-parametric cluster-based permutation tests (CBPTs) (Oostenveld et al., 2011), including all channels, time points, and frequencies. We tested the main effect of *Condition* ([Biperiden Learned + Biperiden Random] vs. [Placebo Learned + Placebo Random]), main effect of *Sequence type* ([Biperiden Learned + Placebo Learned] vs. [Biperiden Random + Placebo Random]), and their interaction ([Biperiden Learned – Placebo Learned] vs. [Biperiden Random – Placebo Random]). The CBPT enabled analysis of the neuronal data without a priori assumptions regarding the location or extent of possible effects. The multiple comparison problem is resolved by applying a single test statistic to clusters instead of individual sampling points. Cluster formation was performed using a dependent samples two-sided t-test, with a p-value threshold of p < 0.025 per side, and adjacency was defined as a minimum of two neighboring channels. A permutation distribution was obtained by pooling the averages per participant, irrespective of condition, then randomly assigning them to two categories using a Monte Carlo simulation. For each of 1000 randomizations, t-tests were applied and clusters determined, with the sum of t-values of the maximum cluster per randomization as the cluster-based test statistic. The p-value was then derived by comparing the uncorrected observed cluster-based test statistic with the permutation distribution and was the proportion of randomizations in which the permuted cluster-based test statistic was larger than the observed cluster-based test statistic. P-values below our critical alpha-level (p = 0.025 per side) were deemed significant.

## Results

One participant was excluded from the analyses due to a lack of responses, leaving N = 29 participants (M = 24.1 years, [22.9 25.2]) for the analyses.

### Behavioral findings

Fatigue levels, perception of whether there was a repeated sequence, and ability to produce the repeated sequence did not differ according to *Condition*.

The mean normalized learning score was greater in the Placebo (M = 0.153, [0.12 0.18]) than the Biperiden (M = 0.091, [0.072 0.11]) condition (F(1,28) = 11.5, p = 0.002, η^2^ = 0.29) (Figure 1c).

The RTs were not all normally distributed. One participant was identified as an outlier, having RTs exceeding three times the interquartile range. Examining the performance of this participant separately, accuracy exceeded 11 of 12 items overall for learned and also for random sequences in both the placebo and biperiden sessions, suggesting correct task performance and hence inclusion in the analysis.

There was no significant four-way interaction. A three-way interaction was observed between *Condition, Sequence type*, and *BMI* (F(1) = 5.26, p = 0.030, η^2^ = 0.16) and correcting for *BMI*, between *Condition* and *Sequence type* (F(1,27) = 4.34, p = 0.047, η^2^ = 0.14). Post hoc pairwise comparisons showed a significantly smaller RT reduction to learned sequences after biperiden (M = 10.06, [-1.73 21.8]) than placebo (M = 27.2, [15.9 38.5]; p = 0.018), with no effect on RT change to random sequences (Biperiden: M = -2.17, [-12.9 8.54]; Placebo: M = -4.28, [-18.9 10.4]; p = 0.79; Figure 2). RT reduction showed a trend towards being greater for Learned than Random after biperiden (p = 0.073), and the RT reduction was significantly greater during Learned than Random after placebo (p = 0.003).

**Figure 2.**
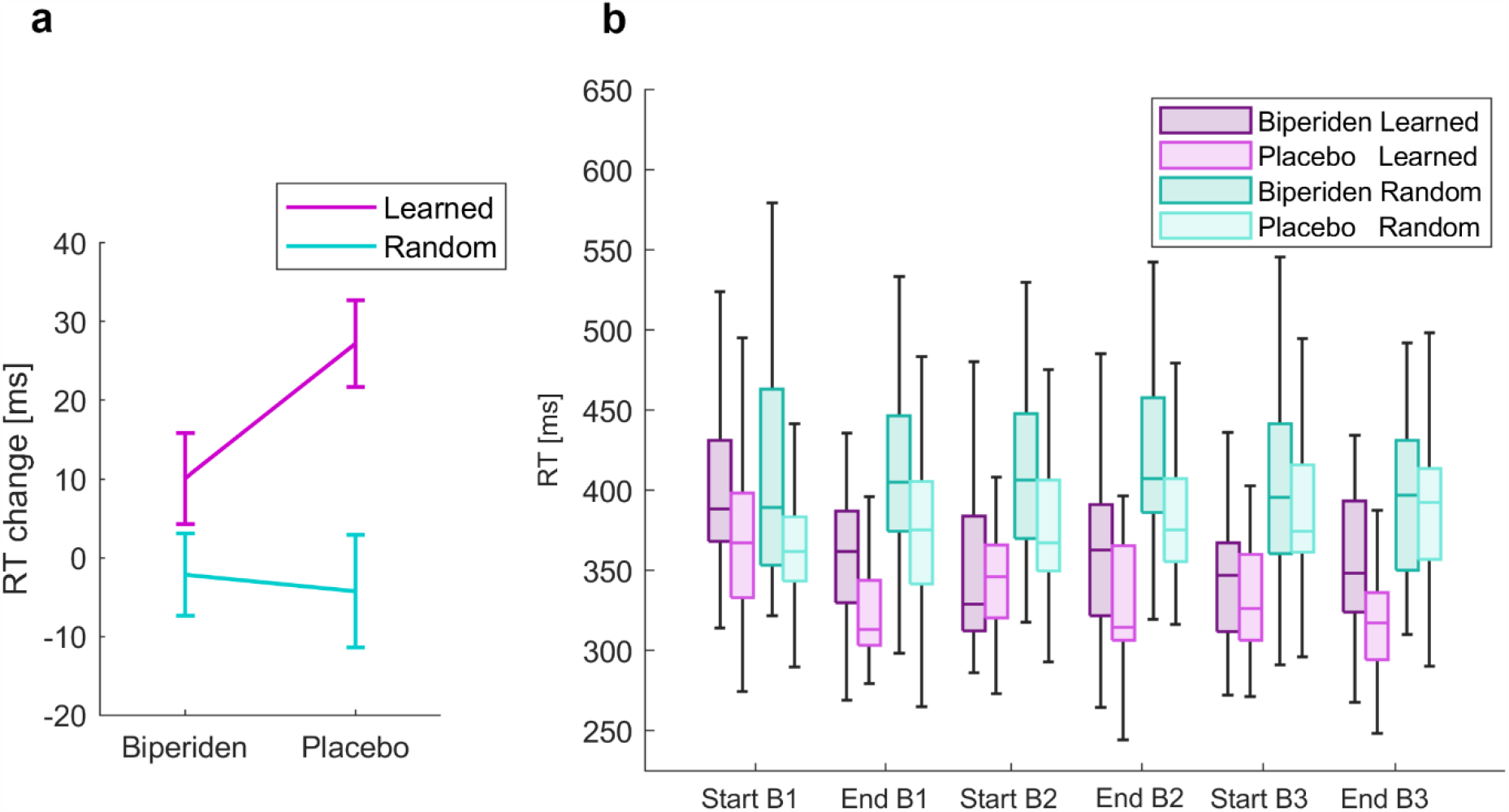
Reaction times (RT). **a**. RT to learned sequences were slower in the biperiden than the placebo condition p = 0.018). RTs to random sequences did not differ between conditions (p = 0.79). **b**. RTs over the course of Block 1 (B1), Block 2 (B2), and Block 3 (B3).

In the biperiden session, no significant interactions were seen between RT changes according to *Sequence type* and either *Session order* or *Time after biperiden given*. Consistent with the main findings, a trend was seen towards an interaction with *BMI* (F(1) = 2.28, p = 0.14, η^2^ = 0.084) and towards RT changes being greater for Learned than Random (F(1,25) = 2.99, p = 0.096, η^2^ = 0.11).

### Electrophysiological findings

The two left-handed participants were excluded from the EEG analysis to enable evaluation of laterality of effects in the remaining participants (n = 27). Grand-average ERSPs were created by averaging over participants (Figure 3). For the interaction between *Condition* and *Sequence type*, we observed a difference in theta, alpha, and beta power (cluster-t = 32,476, p_pos_ = 0.021, SD = 0.005). The cluster was observed over the ipsilateral motor and bilateral parietal–occipital areas, although more pronounced over the ipsilateral hemisphere (Figure 4a).

**Figure 3.**
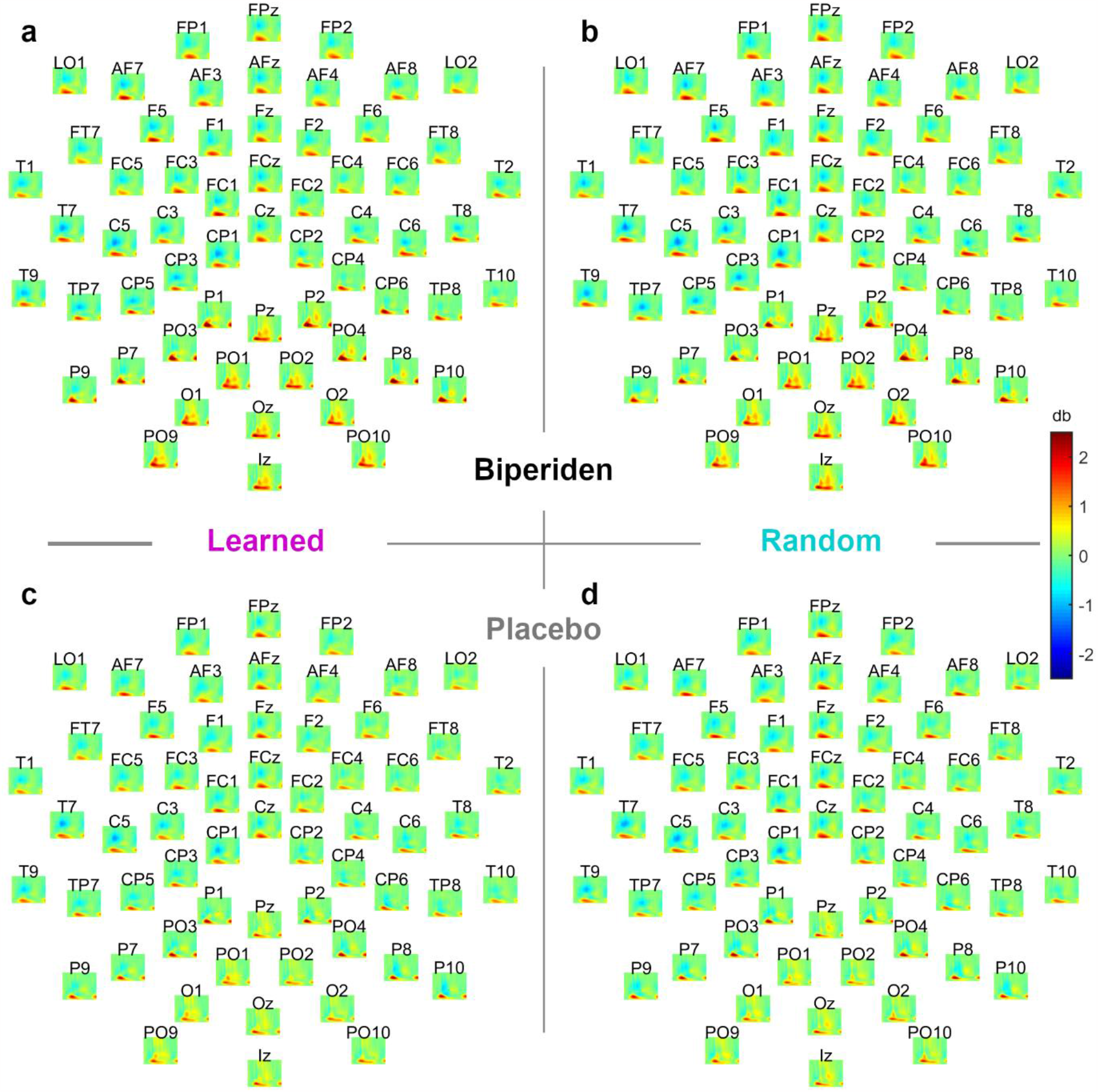
Event-related spectral perturbations. **a**. During Learned with Biperiden. **b** During Random with Biperiden. **c**. During Learned with Placebo. **d**. During Random with Placebo. x-axis: -200 ms to 1200 ms; y-axis: 2-40 Hz. Colorbar applies to all subpanels.

**Figure 4.**
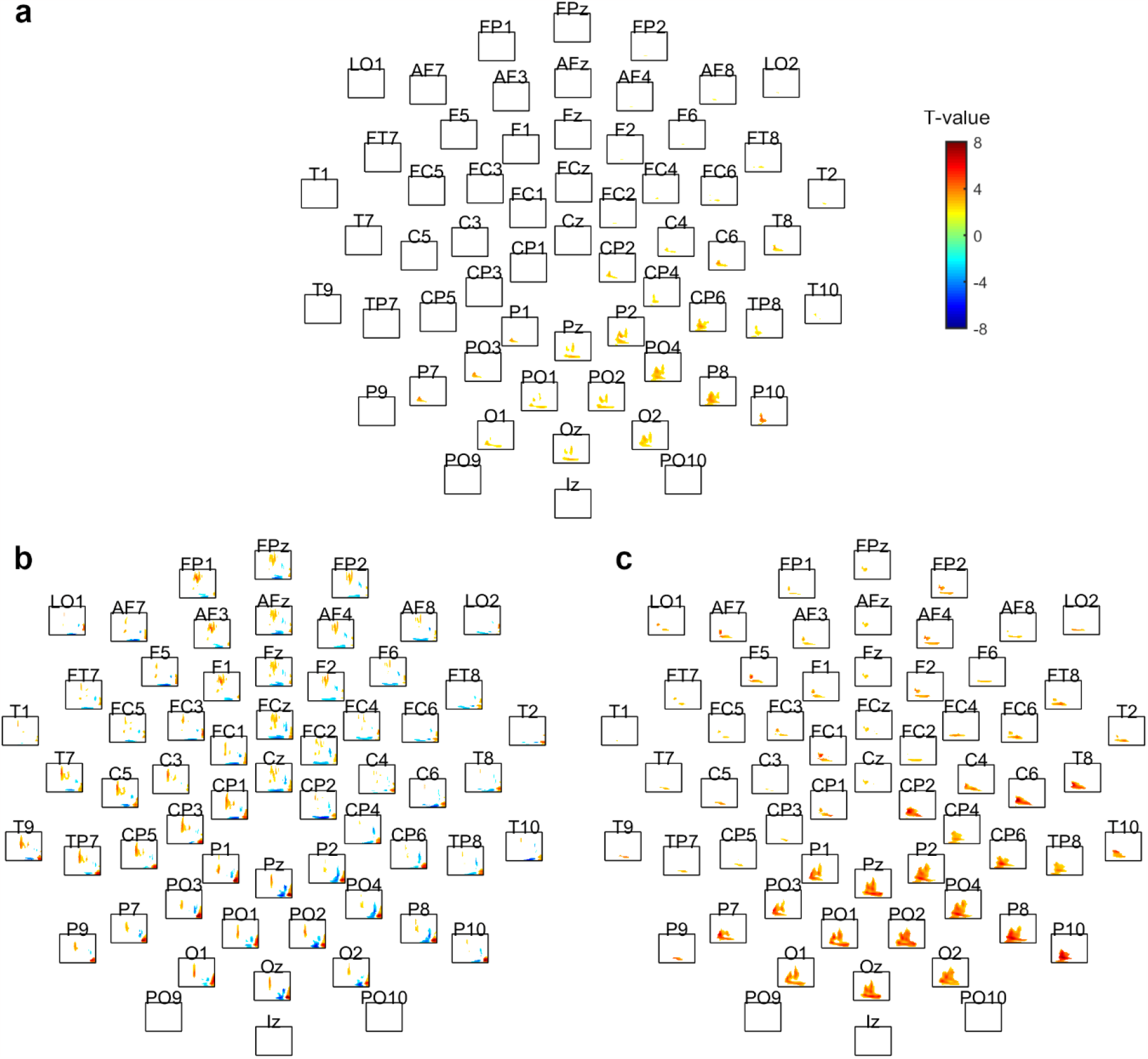
T-value maps. **a**. Interaction between *Condition* (Biperiden, Placebo) and *Sequence type* (Learned, Random). **b**. Main effect of *Sequence type*. **c**. Main effect of *Condition*. x-axis: -200 ms to 1200 ms; y-axis: 2-40 Hz. Colorbar applies to all subpanels.

For the main effect of *Sequence type*, we observed a difference between Learned and Random in three clusters (Figure 4b). Cluster 1 showed an earlier and greater theta and alpha power during Learned compared to Random around 1 s post-stimulus (cluster-t = 54,408, p_pos_ = 0.014, SD = 0.004). The cluster included a widespread area but was most pronounced fronto–centrally and over a bilateral parietal–occipital area. Cluster 2 showed less and shorter beta power desynchronization during Learned than Random (cluster-t = 47,704, p_pos_ = 0.014, SD = 0.004). The cluster was widespread over a central frontal and a bilateral centro–parietal–occipital area, with some lateralization to the contralateral hemisphere. Cluster 3 showed lower theta power peaking at around 400 ms during Learned compared to Random (cluster-t = -64,390, p_neg_ = 0.004, SD = 0.002). The cluster was widespread. Over the parietal–occipital area, it additionally encompassed the alpha and lower beta band.

For the main effect of *Condition*, Biperiden and Placebo differed (cluster-t = 144,381, p_pos_ = 0.002, SD = 0.001). Theta, alpha, and beta power were greater after biperiden than placebo (Figure 4c). The main part of the cluster incorporated a bilateral parietal–occipital area starting shortly after stimulus onset and persisting until 600 ms post-stimulus onset. Theta power differences were observed widespread bilaterally. Alpha power differences were more lateral over the ipsilateral motor area and a bilateral parietal–occipital area. Beta power differences occurred mainly over a bilateral parietal–occipital area.

## Discussion

Biperiden administration was associated with a specific impairment of motor sequence learning in a healthy participant group. Normalized learning scores were lower after receiving biperiden than after a placebo. A two-way interaction was seen for the RTs between whether biperiden or a placebo was given and whether the sequence was learned or random. Motor sequence learning performance was impaired following biperiden compared with after placebo, as reflected by a smaller RT reduction to a repeated sequence, while visuomotor responses to random sequences did not significantly differ according to condition. The absence of an impact of biperiden on performance during random sequences suggests a specific modulation of sequence learning rather than general motor execution and fits with previous studies detecting no impact of biperiden on general motor performance (Borghans et al., 2020; Guthrie et al., 2000), while ACh inhibition impaired consolidation following procedural learning (Rasch et al., 2009).

An interaction between *Condition* and *Sequence type* was also observed in the effects of cholinergic modulation on oscillatory power. We consider this interaction in the context of known oscillatory correlates of sequence learning and of cholinergic modulation.

The oscillatory power differences between motor sequence learning and responses to a random sequence are in accord with previous studies. We observed widespread lower theta power peaking at around 400 ms (Cluster 3) and earlier and greater posterior theta and alpha power around 1 s post-stimulus (Cluster 1) during learned compared to random trials. Fronto–central theta power is modulated by the attentional demands associated with processing and responding to sensory information (Clayton et al., 2015; Lum et al., 2022; Sauseng et al., 2007; Tzvi et al., 2016). An early theta decrease during learned compared to random trials was also observed by (Lum et al., 2022), who suggested that theta desynchronisation reflects top-down attentional mechanisms disengaging from sensory processes, promoting implicit sequence learning. Greater posterior theta and alpha power during a fixed than random sequence is also thought to reflect visuospatial attention mechanisms. Greater theta power during the repeated sequence is thought to reflect working memory processes during sequence learning (Jensen & Mazaheri, 2010; Roux & Uhlhaas, 2014; Tzvi et al., 2016). The post-stimulus theta decrease is therefore consistent with less attentional demand, due to knowlege regarding the sequence, and thus successful learning. The alpha rhythm is implicated in sensory gating (Klimesch et al., 2006), and lower alpha power during stimulus-response mapping for visuomotor responses to the random sequence is associated with greater visual attention.

Also consistent with previous findings is the reduced and briefer beta power desynchronization during learned compared to random sequences over the central frontal, contralateral somatosensory, and contralateral parietal cortex (Cluster 2). Cortical reorganization during sequence and stimulus–response learning is associated with modulation of beta oscillations (Lum et al., 2022; Pollok et al., 2014). Shorter periods of beta desynchronization are associated with better predictability of the appearance of the next stimulus (Alegre et al., 2006), and prolonged periods of beta desynchronization are thought to maintain the current motor state, thus hindering behavioral flexibility (Engel & Fries, 2010; Meissner et al., 2019; Voegtle et al., 2023). Beta oscillations contribute to motor sequence learning by integrating visual and somatosensory inputs (Hardwick et al., 2013; Hazeltine et al., 1997; Voegtle et al., 2023). The shorter desynchronization in contralateral parietal beta power observed during learned trials may facilitate predictability, and therefore information integration, between premotor and parietal areas as a sequence is learned through repetition.

ACh inhibition through biperiden resulted in increased theta, alpha, and beta power compared to placebo. While the theta and beta power increase fits with previous findings (Knott et al., 1997; Osipova et al., 2003; Sannita et al., 1987), ACh inhibition is typically accompanied by an alpha power decrease (Neufeld et al., 1994; Sannita et al., 1987). A reduction in alpha power ratio between eyes closed and eyes open, due to greater alpha synchronization while eyes were open, has been reported, however (Osipova et al., 2003). We note, however, that these studies examined the effects of cholinergic modulation at rest, without attentional engagement. A recent oculomotor theory proposed a link between alpha modulation and saccadic eye movements, even without visual input, as reduced oculomotor activity, which yields a steady gaze, suppressing non-foveal input, is linked with increased alpha power (Popov et al., 2021). Greater alpha power during learned than random sequences would fit with lower oculomotor activity when responses are less dependent on visual input. Biperiden could excessively reduce already lower oculomotor activity, given that participants responded to visual cues, even during learned sequences. Increases in all three frequency bands have also been observed after cholinergic inhibition during movement and navigation in animal studies (Dimpfel, 2005; Dringenberg & Zalan, 1999).

For the main effect of *Condition*, and for the interaction between *Condition* and *Sequence type* we observed the greatest effect over a parietal–occipital area. Cortical changes have previously been reported in a similar area following pharmacological modulation of the cholinergic system (Bauer et al., 2012; Sannita et al., 1987). ACh regulates spatial integration in the human visual cortex (Silver et al., 2008) and modulates spatial attention in the occipital, parietal, and motor cortex through an impact on alpha and beta oscillations, which are known to play a key role in gating sensory processing (Bauer et al., 2012). ACh antagonists are known to affect these visual brain areas and reduce attentional modulation (Danielmeier et al., 2015; Herrero et al., 2008). Biperiden specifically influences neural correlates of prediction and discrimination, suggesting an effect on early perceptual processing (Klinkenberg et al., 2012). It is plausible that the large-scale synchronization induced by biperiden hinders the selective attention required for sequence learning.

Oscillatory power differences following biperiden compared to placebo were more pronounced ipsilaterally. Inclusion of only right-handed participants suggests cholinergic modulation of motor sequence learning rather than motor execution related processes. This asymmetry might be linked to hemispheric specialization (Schubotz & Cramon, 2003; Voegtle et al., 2023) or differential distribution of M1 receptors within the cortex (Disney et al., 2006; Yamasaki et al., 2010).

These findings support the hypothesis that the cholinergic system is engaged in motor sequence learning, with important implications for its interruption with standard movement disorder medication in patients who may already have motor sequence learning impairment. A limitation of the current study is the inclusion only of male participants, due to potential risk in cases of unknown pregnancy. Future work is required to examine whether the deterioration in motor sequence learning is also observed in patient groups taking biperiden, and in older participant groups, in whom pregnancy is no longer a possibility, both female and male participants should be included.

## Conclusion

We provide evidence that the muscarinic ACh receptor antagonist biperiden has selective effects on cognition. Biperiden specifically impaired motor sequence learning but not general motor execution. Together with modulation of electrophysiological activity in nodes of the motor learning network, the findings provide direct evidence for the engagement of ACh in motor sequence learning.

